# SurfGraphPro: Integrating Protein Language Models with Geometric Deep Learning On Coarse Protein Surfaces for Binding Site Prediction

**DOI:** 10.64898/2026.09.17.751511

**Authors:** Adela Habib, Li-wei Hung, Martha Dix, Kaetlyn Gibson, S. Chain Patrick, Bin Hu

## Abstract

Accurate and fast prediction of protein binding sites remains essential for understanding molecular interactions and facilitating protein engineering. Here, we present SurfGraphPro, a geometric deep learning approach that bridges protein language models with surface-based structural representations for a rapid binding interface prediction. Our method operates on coarse-graphed triangulated protein surfaces, utilizing solvent-excluded surface meshes downsampled into amino acid residue centered patches. Unlike existing approaches that rely on extensive physicochemical feature engineering, we leverage evolutionary information directly from protein language model embeddings, eliminating the need for computationally expensive multiple sequence alignments and hand-crafted features. This integration achieves on average ***∼* 18 − 28×** speedup for proteins of ***∼***100 to a few 1000s of amino acids over current state-of-the-art surface-based model while maintaining comparable accuracy on diverse binding interfaces, including challenging antibody-antigen complexes. To our knowledge, this represents the first approach to integrate protein language model embeddings with coarse geometric surface representations for binding site prediction, demonstrating that learned evolutionary features coupled with geometric transformers can replace traditional feature engineering without sacrificing performance.

## 1 Main

Protein-protein interactions (PPIs) form the foundation of cellular processes, from signal transduction to immune response and metabolism Keskin et al. (2008). The ability to accurately predict protein binding sites has profound implications for drug discovery, protein engineering, and understanding disease mechanisms Gonzalez and Liao (2012). Structural-based prediction of binding interfaces offers precise insights into the physical basis of these interactions, with protein surfaces representing the critical interface where molecular recognition occurs.

Predicting binding interfaces using protein surface presents several challenges. First, protein surfaces exhibit topological complexity with binding interfaces often spanning discontinuous amino acid sequences. Second, binding interfaces vary greatly in characteristics-from deep pockets in enzyme-substrate complexes to flat surfaces in antibody-antigen interactions Jones and Thornton (2012). Third, the high dimensionality and feature complexity of protein surfaces make computational approaches challenging.

Early computational methods on protein surfaces included geometric hashing Fischer et al. (1995), patching analysis Jones and Thornton (1997), electrostatic complementarity analysis McCoy et al. (1997), and conservation-based scoring Lichtarge et al. (1996). These approaches relied on Connolly’s molecular surface algorithm Connolly (1983) and deterministic feature extraction, which is often computationally expansive. Machine learning subsequently transformed this field with random forests Qi et al. (2005), convolutional neural networks (CNNs) on voxevelized proteins Jiménez et al. (2018), and graph neural networks (GNNs) modeling residues as nodes Fout et al. (2017). Protein language models (PLMs) have also been applied in PPI predictions, using separated protein embeddings Sledzieski et al. (2021, 2023) or longer context to handle a pair of protein sequence directly Liu et al. (2025). However, recent studies showed that PLMs are a source of data leakage for PPI predictions due to the inherent strong memory of cross-attention networks, capturing shared evolutionary history Szymborski and Emad (2026). When data leakage is avoided by minimizing sequence similarities between training and test datasets, PPI prediction performance could drop significantly Park and Marcotte (2012); Bernett et al. (2024).

Among those explored machine learning methods, geometric deep learning approaches have shown particular promise. MaSIF Gainza et al. (2020) pioneered geodesic CNNs on protein surface meshes using sophisticated feature engineering. Other notable approaches include Docking-GNN Yuan et al. (2024) employing GNNs with attention mechanisms, ScanNet using multiple sequence alignment (MSA) with attention networks on atomic and residue graphs Tubiana et al. (2022), PeSTo applying geometric transformers directly on protein heavy atom graphs Krapp et al. (2023), DeepRank-GNN Ŕeau et al. (2023) combining graph representations with physics-based potentials, and DeepSurf Mylonas et al. (2021) implementing surface-based CNNs with multi-scale geometric features.

Despite these advances, current methods require extensive feature engineering Gainza et al. (2020); Mylonas et al. (2021); Tubiana et al. (2022); Xu et al. (2024) or depend on physics-based scoring functions that may not generalize well across diverse protein families Ŕeau et al. (2023). Additionally, several approaches struggle with computational efficiency when analyzing large protein complexes beyond 1000 amino acid residues Jiménez et al. (2018); Krapp et al. (2023).

In this work, we present SurfGraphPro, which makes three key contributions to protein binding site prediction modeling. First, we demonstrate that protein language model embeddings (ESM-2, Lin et al. (2023)) can effectively replace hand-crafted physicochemical features while capturing evolutionary information without requiring multiple sequence alignments or atomistic property calculation. Second, we introduce an efficient surface representation through residue-centered patch-based coarse-graphing of triangulated molecular surfaces, enabling scalable geometric deep learning on protein surfaces. Third, we bridge these two paradigms—language models and geometric learning—through E(n)-equivariant geometric transformers that preserve physical symmetries. This approach not only achieves performance comparable to state-of-the-art methods with significantly reduced computational cost ( 18-28 ) on large proteins but also provides a generalizable framework for integrating sequence and structure information in protein analysis.

## 2 Results

### 2.1 SurfGraphPro model architecture

SurfGraphPro as shown in Fig. 1 takes in coarse graphs of protein surfaces and returns binding probabilities of surface vertex points and surface amino acids in two modes: (1) partner-agnostic, Fig. 1 (b); and (2) partner-specific, Fig. 1 (c). In the partner-agnostic case, the model can identify all regions on the surface as binding or not regardless of who those binding sites belong to, see Fig. 1 (b). In the case of partner-specific mode, when two protein surfaces, a target and its partner, are concerned, SurfGraphPro as shown in Fig 1 (c) applies bi-directional cross-attention mechanism to predict if the two proteins binds and where they would bind on their surfaces.

**Fig. 1:**
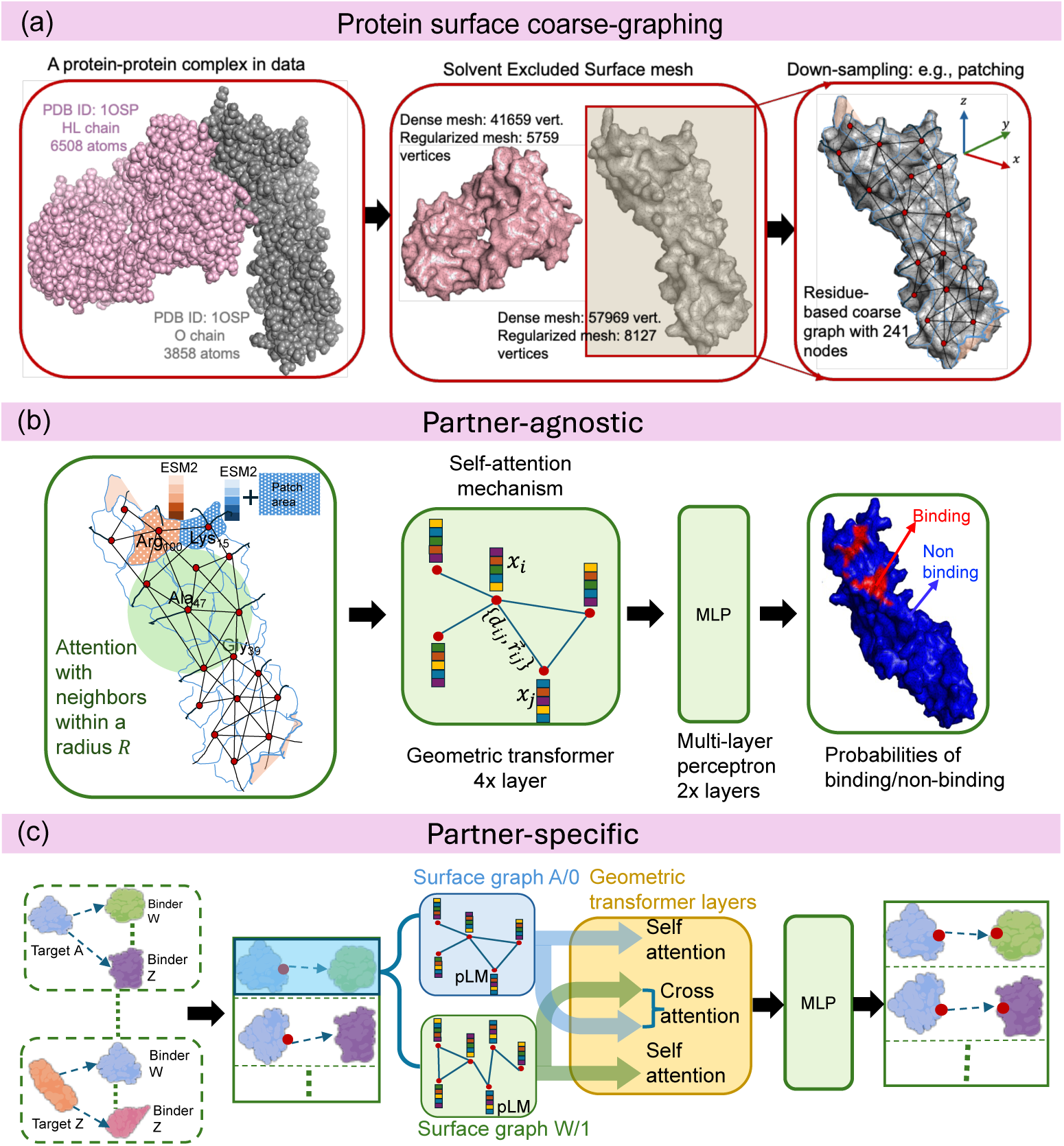
A schematic of SurfGraphPro model architectures. used for partner-agnostic and partner-specific binding site predictions. (a) Given 3D structures of a protein complex, we build 3D triangular meshes which then gets converted into a coarse surface graph. (b) The partner-agnostic model comprises of four geometric self-attention layers, where each nodes attend to all the neighbor nodes within a radius of *R*. Subsequently, a two layer multi-perception block decodes the residue-surface graph node outputs into probabilities of 1 (binding) or 0 (non-binding). (c) In the partner-specific model, inputs to the model is two coarse residue-based surface graphs of a target and a binder. In addition to self-attention layers, we have cross-attention weights that also update the node features within the four geometric transformer layers. Here too, two multi-layer perceptrons decodes each nodes context information for each of the proteins in the pairs into binding/non-binding probabilities.

### 2.2 Partner-agnostic binding site predictions

Model outputs are probabilities that determine if a surface-exposed amino acid residue is binding or not. So, for each protein chain in our Test 358, we get per protein per surface residue predictions. Figure 2 presents metrics from evaluating SurfGraphPro on the Test 358, along with benchmark tests done on a subset of 53 protein chains that represent transient interactions, demonstrated to be difficult to capture in most AI/ML binding prediction models.

**Fig. 2:**
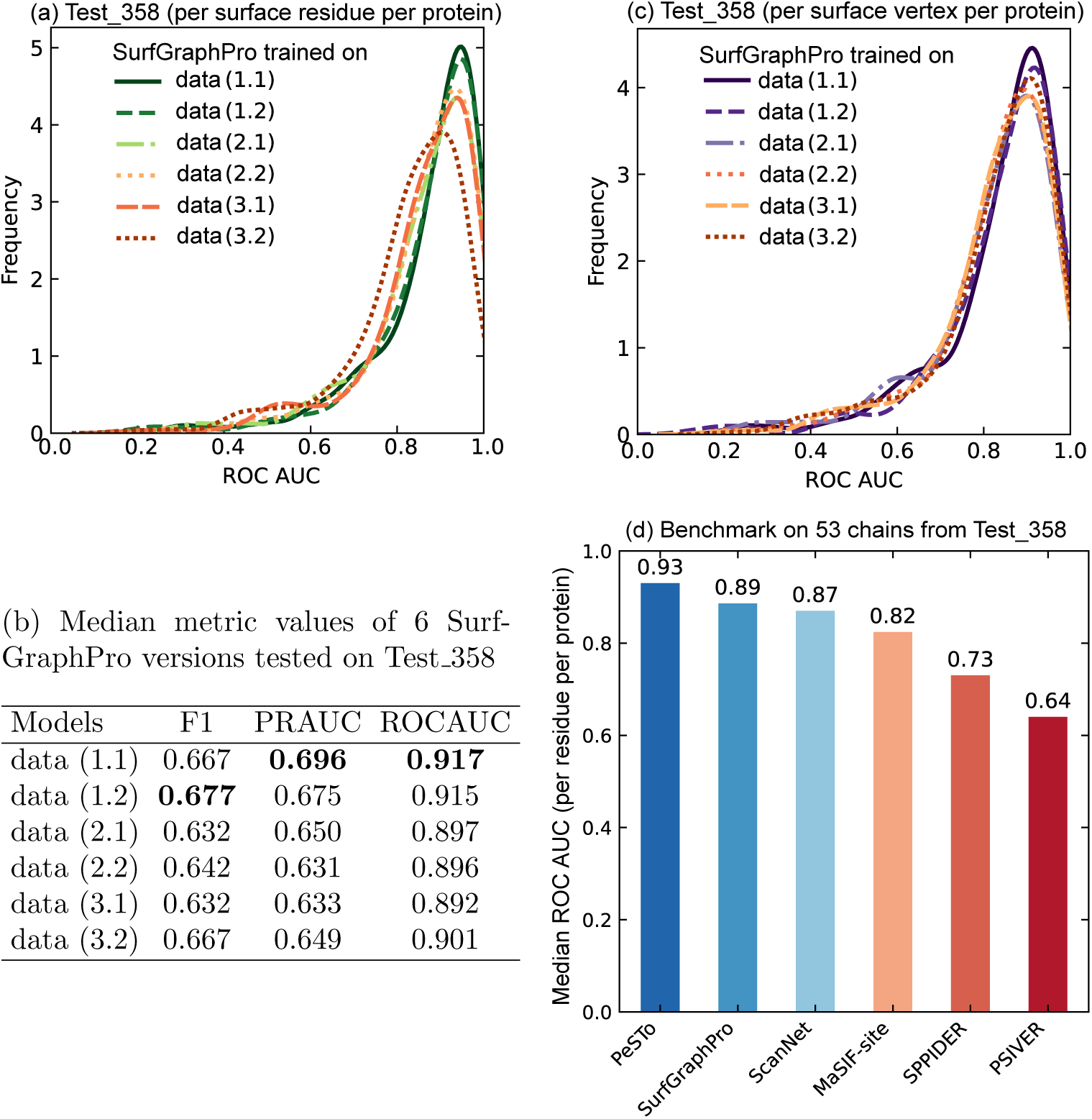
Partner-agnostic models’ predictions on Test 358. Histogram density plots of ROC AUC metric for the six SurfGraphPro model versions trained on six data sets with varying degree of redundancy reduction and data leakage checks. (a, b) present ROC AUC computed for each protein in the Test 358 set that include all types of interactions including polar/non-polar, hydrophobic/hydrophilic. Panel (a) shows predictions done per residue on the surface and (c) shows predictions per surface vertex points. Table in panel (b) shows median values of other metrics such as F1 and PR AUC for these SurfGraphPro model versions tested on Test 358. Panel (d) shows comparison of median ROC AUC per protein at per residue prediction of SurfGraphPro with existing state-of-the-art binding-site prediction models, from fully-atomistic model of PeSTo to the purely sequence-based PSIVER model. The results in panel (d) reveal that coarse-graphing places our model in between the fully atomistic PeSTo model (performance smaller than by *∼*4.5%) and the fully surface-vertex-based MaSIF-site model (performance greater than by *∼*8.5%).

First, in Fig. 2 (a) we show how six different versions of SurfGraphPro perform when tested on the Test 358. These six different versions of the SurfGraphPro are trained on six datasets derived from the original PPI list from the Dockground database after applying two levels of data leakage and data redundancy reduction approaches, see section 4.1.1 for details. The data redundancy reduction approaches check for any structural and/or sequence level similarities between any protein in the training dataset with all other samples in the training dataset, resulting in three derivatives from the original list of PPIs. The data leakage approach then takes all proteins in these three redundancy-reduced subsets and further checks if there is any exact sequence similarity of ≥30% between these proteins and any proteins in the Test 358, making subsets of data (1.1), data (2.1) and data (3.1). A subsequent step of data leakage analysis also checks if any associated chains in the training data protein complex have any sequence similarities of ≥ 30% with any proteins in Test 358, resulting in subsets of data (1.2), data (2.2) and data (3.2). The goal of this analysis is to check if any redundancy via sequence/structural similarities in training datasets impact the model learning and if any possible data leakage between the test and train sets exist. As seen from the histogram density curves of ROC AUC on Test 358 in Fig. 2 (a), the peaks shift to the left as the training data is reduced by removing redundant samples from training data (see differences between data (1.1), data (2.1) and data (3.1) models). Removing associated protein chains from training data because they share ≥ 30% sequence similarity with samples in the test set does not impact the performance much (see differences between data (1.2), data (2.2) and data (3.2)). These two observation make two points: (1) shifts from redundancy removal could be a result of losing surface-level diversities in the training data. If samples within the training data share structure and/or sequence similarities with other data points in the same training data, that does not necessarily mean their 3D surfaces will also be very similar. Binding pocket geometries can hold various curvature and shape forms which are local attributes and cannot be considered similar when two protein structures show high 3D similarities. (2) Removing samples from train data that share exact similarities with samples in the test set already provide a robust way of data leakage prevention between testing and training. Table 2 (b) presents more median metric values including the PR AUC and F1 to further corroborate how filtering training dataset for redundancy impact the model performance. Comparing PR AUC values from models trained on data (1.1), data (2.1) and data (3.1), we see a reduction of ~10% in accuracy. Importantly, note that in Table 2 (v), median metrics of F1, PR AUC and AUC ROC are all above 0.5, indicating the models’ ability to learn important signatures of binding. Our models are inherently encoding 3D surface level information. However, we coarsen the surfaces to enable rapid and scalable learning of binding sites. This coarsening then prompts the question of how models trained on various training data sets do when the predictions are mapped from the coarse-graphs back to the whole surface. Figure 2 (b) shows histogram densities of ROC AUC from predictions per surface vertices on the proteins in Test 358. Comparing residue level predictions in panel (a) to the surface-vertex level predictions in panel (b), all models follow similar trend aross all proteins in Test 358 with some slight variations, implying that coarse-graphing does not result in any major loss of accuracy when compared with finer-mesh binding site predictions. For models trained on heavily filtered redundant training data, coarse-graph label predictions show more drop in peak density value vs. when predictions are mapped on to the finer vertex-level. For example, model of data (3.2) seems to perform relatively worse than the other models when predicting per surface residues in panel (a). However, per surface vertex point predictions of this same model in panel (a) follow similar trend and is not far off from the rest of models’ predictions. This indicates that even low-performing models trained on coarsened surfaces can capture binding pockets on the actual fine surface meshes relatively well.

Since SurfGraphPro model outputs are per surface residue probabilities, this also allows us to compare our models’ performance against other existing protein binding site identification models which include SPPIDER Zhou et al. (2006), PSIVER Murakami and Mizuguchi (2010), MaSIF-site Gainza et al. (2020), ScanNet Tubiana et al. (2022) and finally the fully atomistic model PeSTo Krapp et al. (2023). Figure 2 (b) shows our SurfGraphPro model’s performance against these models. The benchmark is conducted on 53 proteins from Test 358 that existing state-of-the-art protein binding models have previously used a benchmarking set. These 53 protein single chains have been identified to participate in transient interactions with complex transient binding sites. These protein examples are considered to be difficult examples for deep learning PPI models as these deep learning models are mostly trained on static protein structures with little transient interaction representation. From median ROC AUC values in Fig. 2 (d), SurfGraphPro’s performance is greater by ~8.5% than the fully surface-based model MaSIF-site but performing a bit lower (by ~4.5%) than the fully atomistic PeSTo Krapp et al. (2023). Performance improvement over MaSIF-site is expected because of SurfGraphPro’s architectural design that considers attention between all surface points at various tensorial ranks (0, 1, 2, etc.) at the same time without radius-based disjoint patching approach that MaSIF-site applied, and pLM embeddings provide the important evolutionary signal. The difference in the performance with PeSTo might originate from using a completely different dataset and labeling method for training. PeSTo was trained on a much larger dataset, *>* 100k, single chains which also included various biological assemblies and “Models” within a PDB structural file. Additionally, SurfGraphPro and PeSTo’s labeling mechanism differs: SurfGraphPro uses surface points to find buried pockets, thereby labeling surface vertices which then get converted into labels of surface exposed residues. PeSTo on the other hand labels any residue that has interatomic distance of 5Å. This kind of labeling can lead to more balance between binding/non-binding labels which is helping when training. So, the comparisons with PeSTo do not strictly consider same number of binding and non-binding points per protein.

### 2.3 Partner-specific predictions

The bi-directional cross-attention mechanism (Equation 8 in section4.5.2) enables each protein’s surface representation to be refined based on its partner’s geometry and chemical features. This bidirectional information flow could improve binding site localization, potentially for interfaces with complex geometric complementarity such as antibody-antigen interactions.

First, in Figure. 3 (a), we evaluate how the partner-specific mode performs on a held-out test PPI Test PPI 45. ROC AUC results are divided into two classes: pairs that are part of homo-oligomer complexes and pairs that belong to hetero-oligomer complexes. The goal is to check how interface shape impacts the model’s ability to capture geometric complementarity. Panel (a) shows that in terms of ROC AUC, the model has a clear preference for homo-oligomers with the median ROC AUC greater than 0.9 as depicted by the kernel density estimate (KDE) curves. For the hetero-oligomers, though the ROC AUCs are not as well concentrated, majority of proteins still show predictions are decent well above values of 0.7. From this test set, it seems that the partner-specific model learns better when samples are homo-oligomers, most likely because of well-defined interface geometries and the partners sharing same structural homology. However, this could also be very well due to under-representation of hetero-oligomer samples in the training dataset.

**Fig. 3:**
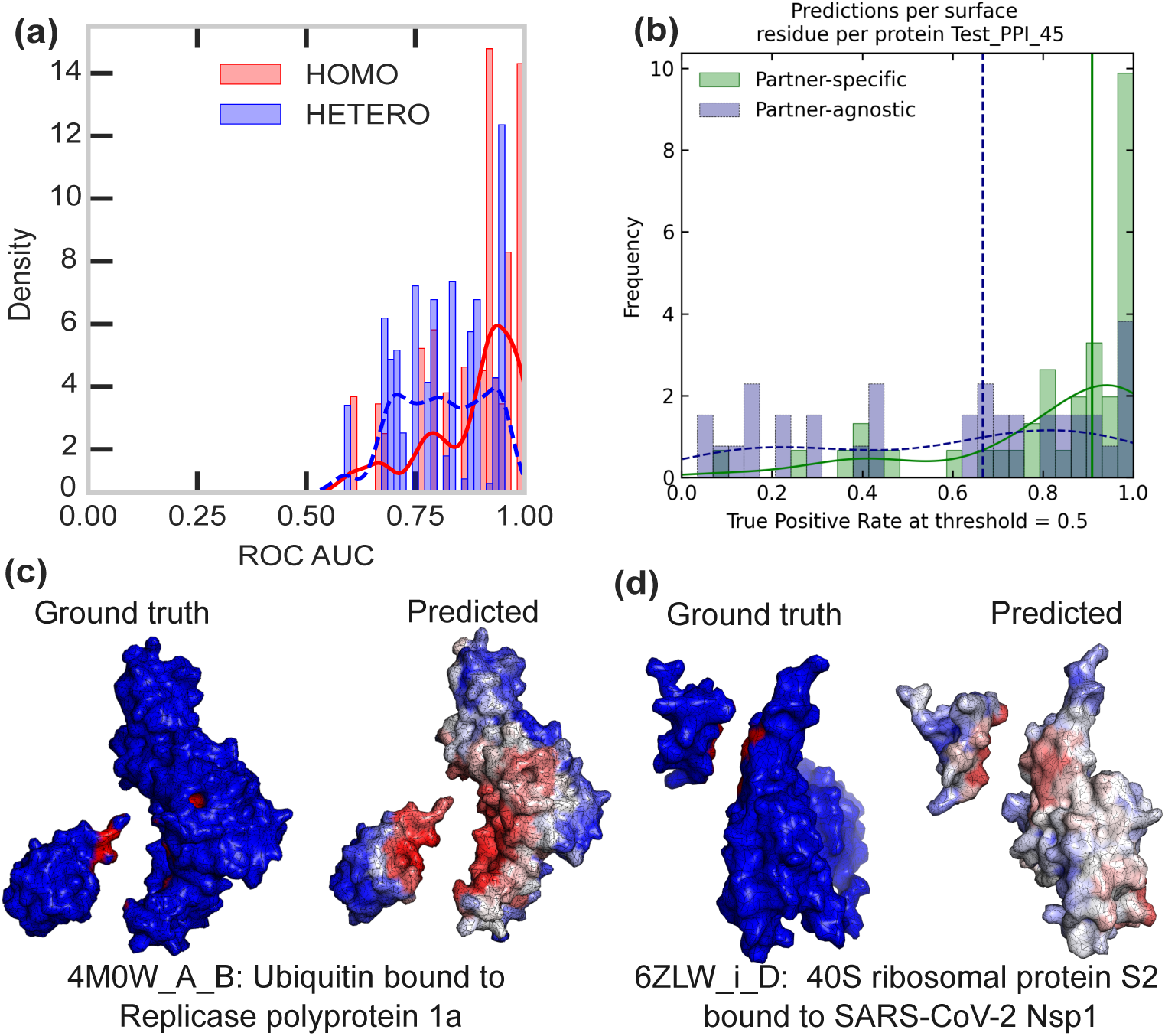
Partner-specific model performance on Test PPI 45. (a) Shows the distribution of ROC AUC for protein pairs that form part of either homo-oligmer or hetero-oligimer complexes in Test PPI 45. In ROC AUC, homo-oligomer cases show more samples towards the higher ranges *>* 0.9 than the hetero-oligomer samples, which are not as concentrated. (b) Histogram of true positive rates or recalls for the subset of proteins in Test 358 whose partners are given in Test PPI 45. Purple dashed lines show how well true positives are predicted for protein of PDBid chain1 in Test 358 while green solid lines show how well the same binding site is predicted when the partner is specified such as PDBid chain1 chainPartner. The vertical lines are median predictions with a clear marking that specifying a partner increases the likelihood of pinpointing the true interface. (c,d) shows 3D surfaces of predicted vs. true binding sites for two cases of a binary complex and a multichain complex, not included in training data.

Next question is how well the partner-specific model of surfGraphPro can localize the binding site on the protein target when it is given a specific partner. To check for this, we have plotted true positive rates calculated as 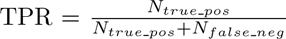 in Figure 3 (b). Here we show all the protein pairs in Test PPI 45 and the same proteins but without their partners from Test 358 using both the partner-specific and partner-agnostic models. Out of 45 cases, 84% of samples have non-zero TPRs when considering the partner-agnostic model from section 4.5.1. In comparison, when partners were specified, 98% of the same protein chains had non zero values of TPRs at the 0.5 accuracy threshold when converting the probabilities to 1. The partner-specific model achieves median TPR of 0.9 compared to 0.66 from the partner-agnostic model on the same protein chains, representing a ~26.6% improvement. This demonstrates that cross-attention mechanisms effectively capture binding complementarity between protein pairs. Figures 3 (c, d) visualize two protein complexes predicted by the model. Panel (c) shows protein complex 4M0W A B: Ubiquitin bound to Replicase polyprotein 1a. This protein pairs belong to the held-out test set of Test PPI 45. The complex is binary and so the only interface is predicted precisely albeit with some broadening on both proteins. Contrary to panel (c), panel (d) shows protein pair 6ZLW i D: 40S ribosomal protein S2 bound to SARS-CoV-2 Nsp1 that is a part of a big complex with many other ribosomal proteins. The model predicts the intended interface between the chains i and D quite well. We notice other binding patches on D as well that from the original structure we know they are binding sites to other partners of chain D. These identified binding sites are considered “false positive” for chain i as they are not specific to chain i. However, could hold a potential/possibility of being a binding site for chain i. This does prompt to a ranking scoring mechanism that is a subject of ongoing work.

Similar to these panels, Figure 4 further visualizes some other cases from the held-out test set how localization of a binding site look like on 3D surfaces compared when no partner is specified. Here, the few examples are chosen of hetero-oligomer proteins that are in Test 358 with no partner while a specific partner is present in Test PPI 45. For most examples where a protein makes interface with a small number of partners, for example 3 partners in Figure 4(a, b), the predictions are able to localize the specific binding site better as is visualized and shown with increased values of ROC AUCs (0.84 vs. 0.92 in panel (a) and 0.84 vs. 0.90 in panel (b)). When the target protein chain makes interface with many other protein chains, i.e., when many binding partners are involved. Figure 4(c) shows protein PDB ID: 3PUZ where 3PUZ B makes interface with 3PUZ F, 3PUZ G and 3PUZ A. When considering just 3PUZ A B, the partner-specific model still finds interfaces of 3PUZ B with 3PUZ F, 3PUZ G in addition to 3PUZ A. Note that ROC AUC = 0.68 reported here takes account of predictions of those “non-specific” partners but the ground truth only accounts for the of the pair input proteins, e.g., 3PUZ A B in here. Overall, it is inferred that when a protein chain has multiple partners, the current model cannot fully localize for just the specific partner given into the model as an input. Future work could improve this localization for a specific partner by adding multiple competing partners through a robust training data sampling or by introduction of some form of bias/gating in the architecture for that specific partner to zero out the competing partners information.

**Fig. 4:**
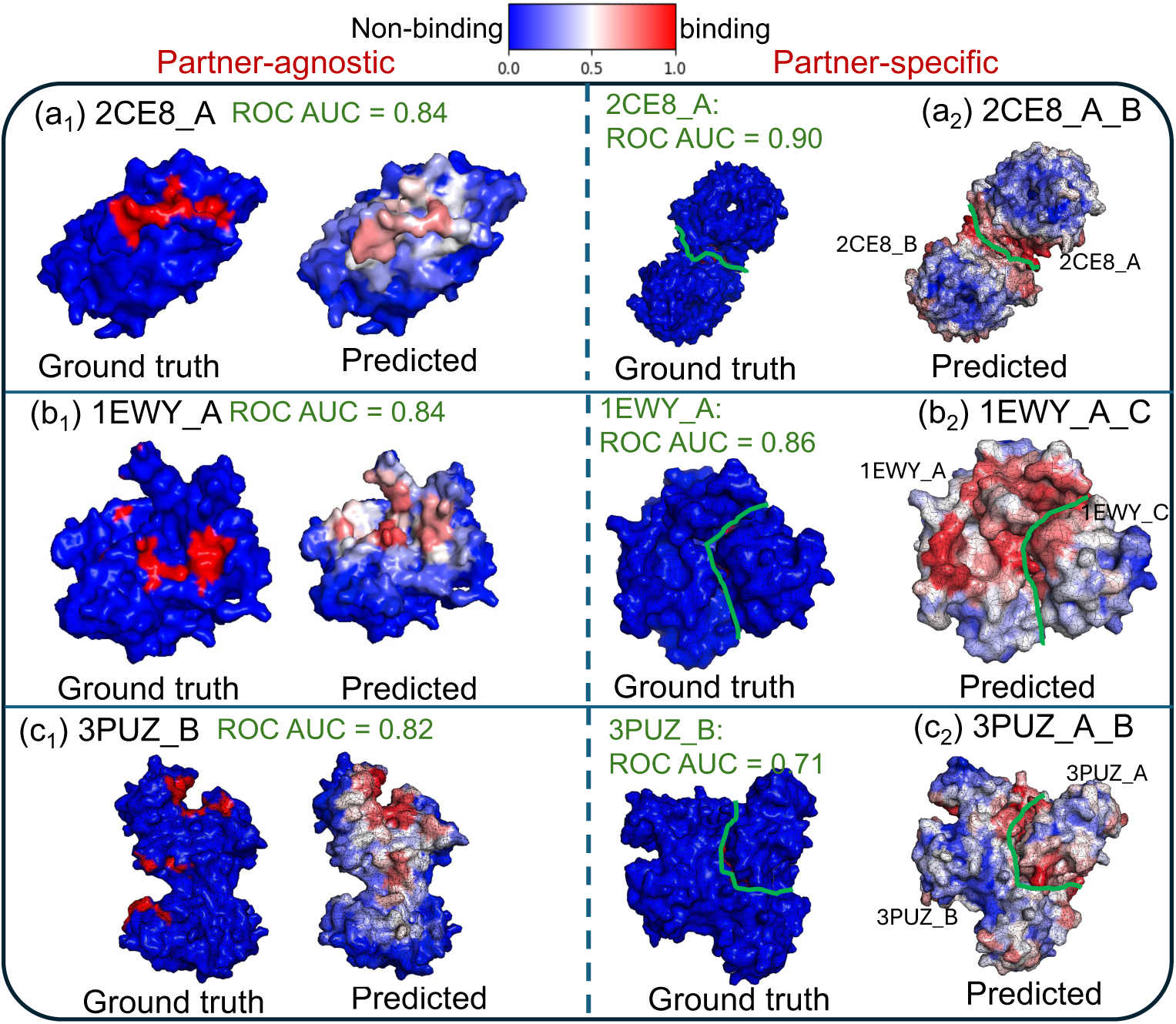
Comparison of partner-agnostic vs. partner-specific on surfaces. Some examples of predictions per surface vertices in the case of partner-agnostic and when the partner is specified. Green lines in column 2 mark interfaces between the two proteins. For panels (a) and (b) ROC AUC increases for the specific binding sites when the partner is given in the model while for panel (c) there is some drop in accuracy. This is most likely because 3PUZ B has multiple partners and the model cannot localized well when only one partner is known.

## 3 Discussion & Conclusion

We present SurfGraphPro, a geometric deep learning framework that bridges protein language models with surface-based structural representations for binding site prediction. By replacing extensive physicochemical feature engineering with protein language model embeddings and the introduction of efficient residue-centered surface coarse-graphing, our approach achieves three key advances: (i) on average 18-28 computational speedup over existing surface-based method, such as MaSIF-site, on protein sizes of 100 to a few 1000s of residues due to elimination of expensive pre-calculation of physicochemical properties and employing coarse-graphing, (ii) elimination of costly multiple sequence alignments while capturing evolutionary information, and (iii) demonstration that learned language model features coupled with geometric transformers can effectively replace hand-crafted descriptors without sacrificing accuracy.

Our residue-centered patch representation exploits the biological organization of binding interfaces while enabling scalable geometric deep learning. The coarse-graphing strategy reduces mesh complexity by 10–100 , making it feasible to train faster on datasets orders of magnitude larger than previous surface-based approaches. In terms of runtimes for binding site identification, Fig. 5 presents comparisons of compute times between MaSIF-site and SurfGraphPro on a GPU and a CPU for 68 proteins of varying sizes (in number of atoms and amino acid residues) sampled from the train data (1.1) in section 4.1.1. While Fig. 5 (a) shows that surface generation task is quite similar in compute time with MaSIf-site with roughly 2× speed gain, likely due to reducing IO overheads, the speed gain from eliminating expensive pre-calculation of physicochemical and geometric features is as large as 1000 for proteins with 3000 residues (Fig. 5 (b)). Note that initially 100 proteins were sampled, but 32 samples were dropped from the figure because feature calculations failed at the APBS Jurrus et al. (2018) level when running MaSIF-site, which further underscores the advantage to avoid pre-calculation of hand-crafted features. At the inference level, Fig. 5 (c) highlights the speed gain of 100 in SurfGraphPro as the protein increases in size because of coarse-graphing in SurfGraphPro. For proteins that have 100-200 amino acid residues as is in Table 5 (d), SurfGraphPro takes 3.47 *±* 0.73 seconds on average using a single CPU and single GPU while MaSIF-site takes 99.50 *±* 17.05 seconds.

**Fig. 5:**
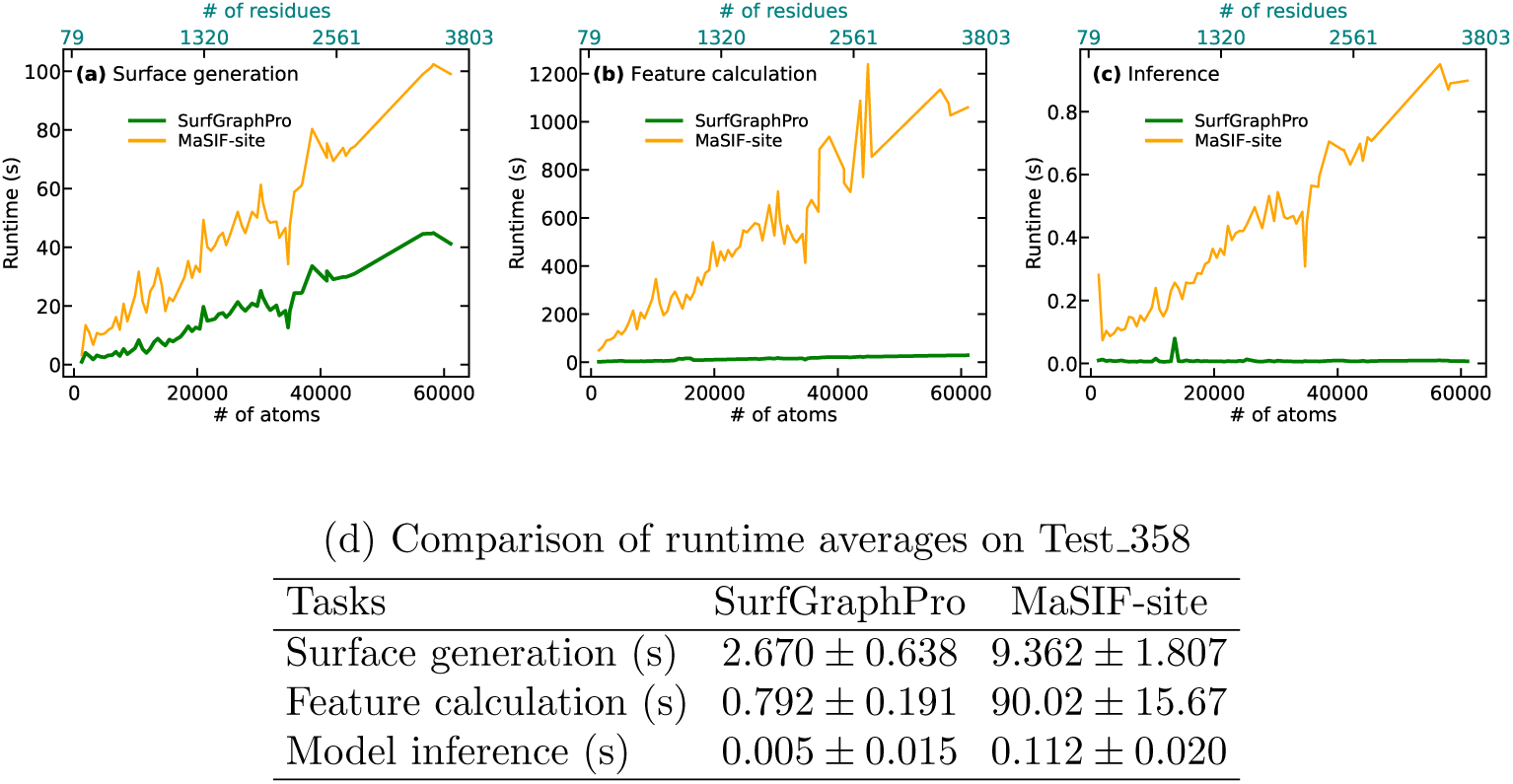
(a) shows the runtime for surface generation that involves running MSMS and pymesh for SurfGraphPro (solid) and MaSIF-site (dotted). Similarly (b) shows the time it takes to do feature calculation which involves physicochemical and geometrical feature extraction for MaSIF-site and for SurfGraphPro that involves surface downsampling, pLM embedding and neighborlist generation using pyTorch. Panel (c) shows inference time for both models using one NVIDIA RTX PRO 6000 Blackwell. (b) implies that by bypassing expensive hand-crafted feature calculations and applying coarse-graphing strategies enables speedup gains up to 3 orders of magnitude. Note that the figure plots runtimes of 68 out of 100 sampled proteins as a function increasing number of atoms (including hydrogens) from train Set (1.1) in section 4.1.1. (d) Table showing runtime averages and standard deviations (seconds) from SurfGraph-Pro and MaSIF-site on 295 protein chains of biological assembly 1 chosen from the Test 358 set.

Currently, the most time-intensive task in SurfGraphPro is the surface generation and mesh processing via MSMS and Pymesh. Both of these steps can be further optimized with advanced meshing algorithms introduced in the field of machine learning for computer vision. For example, dMaSIF Sverrisson et al. (2021) as a successor to MaSIF-site has employed 3D point clouds in PyKeOps Charlier et al. (2021) on GPUs, reducing computation time down to an average of ~ 0.5 seconds for proteins in Table 5. The average inference time of dMaSIF is reported to take 10s of milliseconds with accuracies similar to MaSIF-site. On the other hand, SurfGraphPro’s inference time is on average a few milliseconds (i.e., 100 faster than MaSIF-site) while providing 8% more accuracy than MaSIF-site. Most importantly, as shown in Fig. 5, SurfGraphPro shines for larger proteins containing a few 1000s of residues, which will likely be true when comparing with dMaSIF in the same protein size regime. This is because SurfGraphPro takes in a coarse-graph of a protein surface which contains far fewer vertexes than mesh-defined surface vertexes used in dMaSIF. Nonetheless, we hypothesize that integration of dMaSIF-like surface generation technique with the SurfGraphPro model architecture would provide further performance improvement in accuracy and time for SurfGraphPro, which will be explored in the future.

When modeling protein geometry, there persists a fundamental trade-off between surface-based encoding and full atomistic representation. While atomistic approaches excel at capturing underlying physicochemical properties through explicit representation of bonding interactions including of hydrogen bonds and electrostatic forces, they face significant challenges in scalability and in balancing local versus global interactions. This representational challenge is well documented in the field of machine-learned force-fields for molecular dynamics simulations, where graph neural networks with message-passing architectures predominantly capture local interactions due to computational constraints, often at the expense of neglecting long-range environmental effects that govern global behavior. Moreover, atomistic PPI models that encode experimental protein structures directly in the model as inputs may face challenges with impurities naturally present in experimental structures such as missing atoms and dangling bonds. By adopting a coarse-graph surface-based representation, SurfGraphPro can circumvent the presence of these physically unrealistic atomic configurations, while maintaining the ability to capture essential local and global geometric and chemical features relevant to protein function at computational speed needed for larger protein complexes. We do note that accounting for only surface residues, SurfGraphPro might miss any impacts of the atoms inside the protein fold onto interaction signatures. Since there is no explicit calculation of all-atom features such as electrostatic charges for SurfGraphPro, the accuracy of the model suggests that the impact of in the protein-fold atoms in the interaction is most likely captured via the pLM embeddings. This model interpretation remains as a subject of future investigation.

By integrating protein language models with geometric representations, SurfGraphPro opens new directions for structure-based protein analysis. While we focused on binding site prediction, the ESM-Surface framework is readily extensible to other surface-based tasks such as protein function prediction, epitope identification, and protein design. Future work will explore incorporating confidence scores from language models, extending to protein-small molecule interactions, and integrating dynamics information to capture binding flexibility.

## 4 Methods

Here we discuss surface graph approach we are taking to learn binding sites. Figure 1 presents the workflow, starting from choosing a protein complex in a curated PPI dataset, generating surface meshes on each of the subunit/chains of the protein, then coarsening the mesh. The coarse graphs are fed as inputs to geometric transformers which can then predict if a mesh vertex belongs to a binding site or not. We provide further details about each components of this workflow in the following sections. Note that prior to generating surface meshes, we protonate the PDB files to add hydrogen. This is done by using the Reduce package Word et al. (1999).

### 4.1 Datasets

#### 4.1.1 Partner-agnostic

Building and benchmarking an ML model requires a rigorous regimen to assess performance and generalization while making sure there is no data leakage between the training data and the held-out test data. However, in protein learning space, building a strict pipeline for leakage detection is substantially challenging given that the vast majority of proteins share similarities either in the sequence and/or structure space and experimentally-determined labels are limited. For example, Hummer et al. (2025) showed that predictions of binding affinity when using structural antibody data can show drastic accuracy drops when considering different sequence identity cutoffs between the training and testing data. Similarly, benchmarking models of binding site identifications using a test set is tricky because of both various labeling schemes these models employ and training datasets share structural and/or sequence identities at various thresholds. Therefore, we lay out a comprehensive training data acquisition, redundancy reduction and data leakage pipeline that check for sequence similarities or structure similarities or both against a held-out test set. The held-out test set is chosen to be the 358 (Test 358) single chain proteins curated by Gainza et al. (2020) for binding site identification. Test 358 includes protein chains that are representative of various interactions including polar/non-polar, hydrophobic/hydrophilic and hydrogen bonding. A subset of 53 chains from Test 358 represent transient interactions proven to be difficult when modeling. The section 2 show results of benchmark done on this set to compare against state-of-the-art binding site identification models. Figure 6 presents the training data acquisition, redundancy reduction within training sets and the pipeline to prevent data leakage between the training and testing data. We start with two lists of PPI from the Dockground database Collins et al. (2022): (1) PPI data that has redundant sequences removed at 30% sequence identity matching via MMSeq2 Steinegger and Söding (2017); (2) PPI data that has redundant structures removed by structure alignment via FoldSeek van Kempen et al. (2024) at TM-score threshold of 0.6. From the list (2), we perform further sequence clustering such that none of the proteins share ≥30% sequence identity with other clusters within the training set, thereby getting list (3). Given these three lists with various redundant samples removed, we then perform data leakage by conducting sequence identity matching of each protein in the test set against each protein in each of these training lists. Each list then narrows down to two sub-lists. For example, in data (1.1) we make sure that for all pairs (PDBid*_i_*, PDBid*_j_*) such that PDBid*_i_ ∈* training data (1.1) and PDBid*_j_ ∈* Test 358 do not share sequence identity *≥* 30%. Train data (1.2) is built by taking a step further such that PDBid*_i_* all other chains of the PDBid do not share ≥ 30% sequence identity. This way we build six training datasets each with various redundancy reduction and data leak-proof measures. Figure 6 reports all the numbers of single chain proteins in each dataset after filtering for failed surface generation of the complex for binding/non-binding labeling purposes. Note all the structures are downloaded from RCSB Protein Data Bank Consortium (2025) of biological assembly 1.

**Fig. 6:**
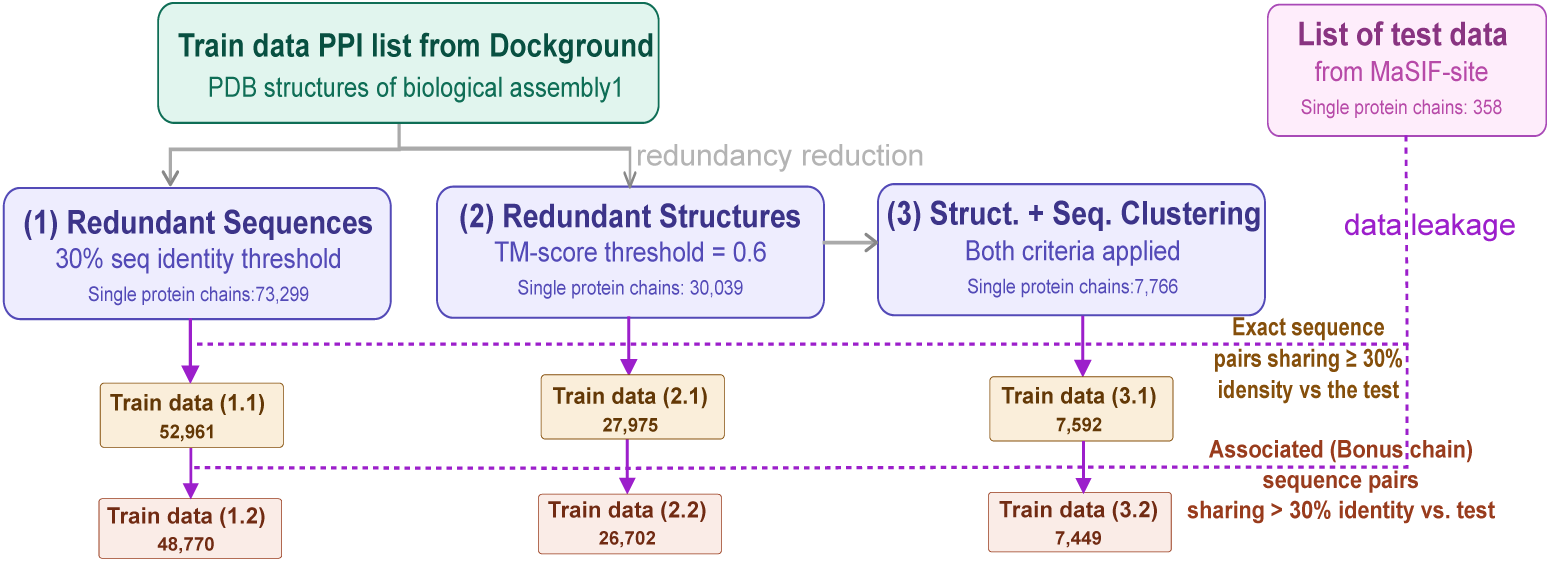
Train and test data curation and data leakage analysis pipeline. Starting with a PPI lists from Dockground database, we download all the PDB structures of the biological assembly one. We take lists from Dockground that have (1) sequence redundancy removed at 30% identity and (2) structure redundancy removed with TM=0.6 score for similarity. On the list (2), we perform further sequence clustering at 30% identity to get list (3). To prevent data leakage, we perform sequence identity matching on these three lists with each protein chains in our benchmark test set from MaSIF-site with 358 single protein chains. This down-selection results in final six training sets at various levels of sequence and/or structure similarities.

#### 4.1.2 Partner-specific

In Figure 6 , list (2) from Dockground database contains 22,262 protein-protein pairs. After filtering for various redundancies and any failed surface generation, we curate a training dataset of 16,569 protein-protein pairs of PDBid chain1 chain2. This list is compiled after removing PDB ids that are present in the held-out test sets of Test 358. For testing purposes, we chose a subset of protein chains named Test PPI 45 such that at least one of the partners is present in Test 358, i.e., if PDBid X is in Test 358, then PDBid X Y is in the Test PPI 45.

### 4.2 Surface mesh creation

For each protein chain in our datasets, we generate a triangulated representation of its solvent-excluded surface at 1 Å resolution using MSMS Imperiali et al. (1993). As defined in Section 4.7, this mesh consists of vertices (3D surface points) connected by triangular faces. The triangulation is subsequently regularized using Pymesh Zhou (2018) to remove degenerate triangles and ensure mesh quality. This fine-grained representation captures the detailed molecular surface geometry but contains too many vertices (typically 10^3^–10^4^) for direct use in geometric transformer training.

### 4.3 Surface mesh downsampling

As described in Section 4.7, we transform the fine-grained surface mesh into a coarse residue-based graph suitable for efficient learning. Our downsampling strategy groups mesh vertices into residue-centered patches. Specifically, for each surface-exposed amino acid, we identify all mesh vertices lying on that residue’s solvent-accessible surface and aggregate them into a single patch. The centroid of these vertices becomes a node in our coarse surface graph , reducing the representation from thousands of vertices to typically 10–1000 nodes per protein chain while preserving the essential geometric and topological information of binding interfaces. See an example shown in Figure 6.

### 4.4 Features and labels

Our coarse surface graph nodes represent surface-exposed amino acids (Section 4.7). We construct node features by concatenating two components: (i) the 320-dimensional ESM-2 embedding Lin et al. (2023) of the residue sequence, which captures evolutionary and biochemical information, and (ii) the patch surface area in Å^2^, computed by summing per-atom solvent-excluded surface areas from MSMS Imperiali et al. (1993). This minimal feature set contrasts with the extensive and computer-intensive hand-crafted descriptors used in MaSIF Gainza et al. (2020), which include hydrophobicity, charge, hydrogen bonding potential, and geometric curvature characteristics of the surface meshes. Note that any non-canonical amino acid found in a data sample is set to letter “X” to be treated that same as is done in ESM-2 tokenizer for unknown amino acids.

Binding/non-binding labels are binary of 1 and 0 respectively. This is done following the method of Gainza et al. (2020), where inter-vertex distances are calculated between the surface mesh of the unbound protein structure and the bound/complex structure. For*_√_*<u>e</u>ach vertex in the unbound structure, if this inter-vertex distance is greater than 2, the empirically known threshold for solvent inaccessibility, then the vertex is assumed to be inaccessible to a solvent molecule, making it a part of the interface upon a protein complex formation.

Since we train on the coarse surface graphs, we need to label the patches whose centers make the nodes. For that we count the number of binding vertices per patch and set a threshold for labeling the whole patch a binding or non-binding. This threshold is set to 5 after systematic evaluation with values of *τ* = 3, 5, 7, and 10, where *τ* = 5 provided optimal balance between precision and recall on validation data (see section 4.6).

### 4.5 Model architecture

As alluded to before, in addition to physicochemical properties, protein-protein interaction is dependent on the local geometry of the surface. Since we are not pre-calculating geometric features, our ML model architecture should be able to perform convolutions on the 3D geometric input data. This leaves us to work with graph-based methods such as graph neural networks or geometric transformers. We choose to work with geometric transformers as this can allow the model to attend to all nodes in the graph and simultaneously restrict edges to neighbors within a radius of choice. In essence, this will be capturing interaction signatures within a local neighborhood while knowing a degree of global environment information. In the sections below, we briefly provide details of the geometric transformer layers used in our models.

#### 4.5.1 Partner-agnostic

Figure 1(b) presents our model architecture for the case of predicting binding sites on a protein when the partner is unknown. This type of prediction is helpful in the case of in-vivo binding for a protein since partners are usually unknown. The input to the model is coarse surface graphs of protein chains with node features from the ESM2 embedding concatenated with the normalized patch areas, 3D coordinates of each vertex nodes, list of neighbors within a user-specified radius of *R*. The model has two blocks, the geometric transformer block to encode and embed interaction signatures and the MLP decoder block for computing probabilities for each node classifying it to binding (1) or non-binding (0).

##### Algorithm 1 Coarse Surface Graph Generation

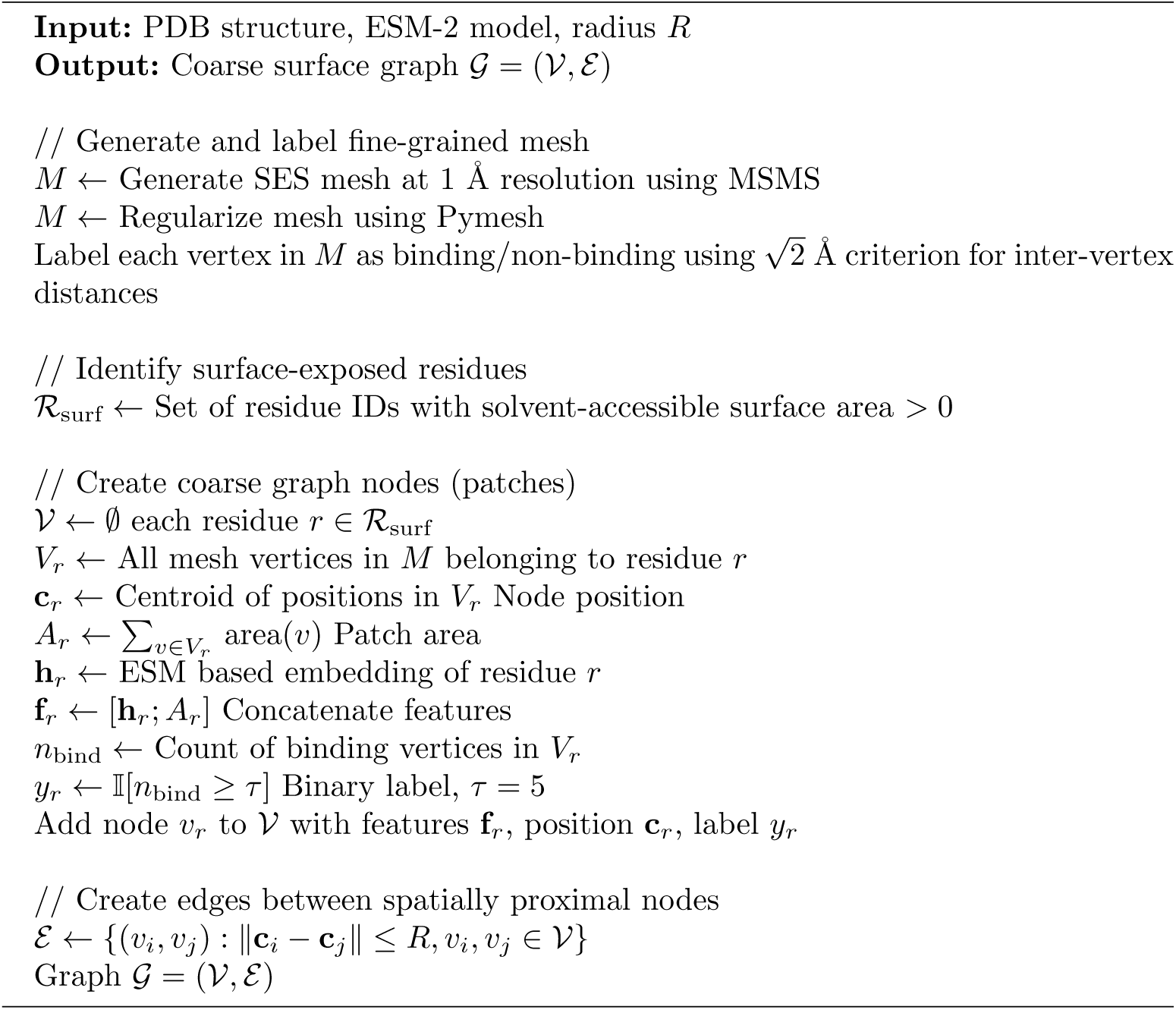

The geometric transformer block takes inspiration from the approach taken by Krapp et al. (2023) for their binding prediction model at the atomistic level with modification relevant to our protein surface coarse-graphs as we discuss here. The geometric transformer block creates a state vector *s* associated with each node *i* in *N* in our graph and refines it throughout the training. This state vector *s* has scalar encoding of *q*∈R*^N×S^* and similarly a vector component ***p***∈R*^N×3×S^*. The dimension *S* is a state encoding dimension specified as a model input. The scalar component *q* is initialized by encoding the node features. We use ESM2 embeddings concatenated with the patch surface area for each node encoded through a multilayer perceptron (MLP) of output size *S*. The vector component ***p*** is learned throughout training and could be initialized as surface normal vectors ***n*** corresponding to each node of the coarse graph at the start of training. For the models reported in this paper, they are initiated as vector of zeros.

A state vector per node is defined as

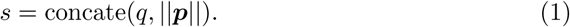

Both the *q* and ***p*** of a node *i* get updated based on encodings of node features and edge features of *i* and all of its neighbors *j* = 0, 1, 2, …, *m*. *j∈nn* with variable size *m* of the neighbor list of *i* within radius *R* defined as a model input. These two features are set as

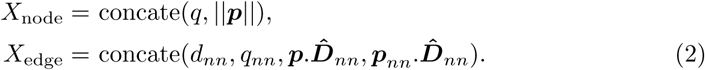

Here, node features are the same as the state vector *s* and the edge features comprise of inter-node distances between *i* and all neighbors *j* of *i* in the set *nn*. The distance is computed as the L2-norm *d_ij_* = *||****r****_i_ −* ***r****_j_||*_2_ and the normalized displacement vector for all *j* neighbors of *i* is defined as 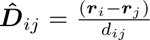 . The *q_nn_* corresponds to the node scalar states of all neighbors *j* of *i ∈ nn*.

Given these node and edge features, we calculate the essential components of self-attention block, i.e., the query, key and value tensors for both *q* and ***p***. The query tensor *Q* is constructed by encoding the node features *X*_node_ using an MLP, which gets the dimensions of *N ×* 2 *× S*, where the 2 corresponds to *q* and *p*. The associated *q* and ***p*** key tensors are encoded by edge features *X*_edge_ through separate MLPs for each, with a distinction that *K****_p_*** has vectorial nature with dimensions of *N × n ×* 3 while the dimension for *K_q_* becomes *N × n*. For the value tensors, we get

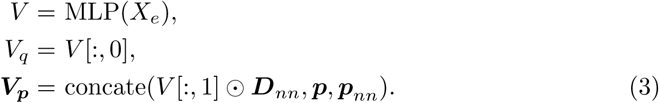

The residual values of *q_h_* and ***p****_h_* are calculated using attention scores from the these query, key and value tensors as

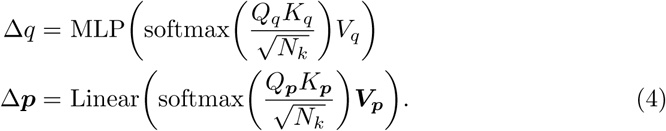

Note that as in original setup by Krapp et al. (2023), the geometric block preserves translational invariance and rotational equivariance. Translation invariance is implied by using 3D coordinate-origin agnostic namely the inter-node distances and displacement vectors. The rotational equivariance is only applicable to vectorial values ***V_p_*** where the only operations on the vector are dot products, scalar multiplications and linear combinations.

Finally, the updated *q* and ***p*** as *q^′^* = *q* + Δ*q* and ***p^′^*** = ***p*** + Δ***p*** update the state vector *s* to *s^′^* = concate(*q^′^* + *||****p^′^****||*). For the binding site classification task, these embedded state vectors get processed through an MLP block to out probabilities for binding/non-binding per node on the surface coarse graph.

The model is trained by minimizing an adaptive-weighted binary cross entropy (bce) loss. The weights were adaptively changed throughout the training steps to account for the imbalance in the number of residue nodes that are binding vs. those that are not binding as

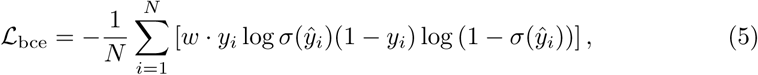

where the weights *w* is calculated for each batch of size *N_b_* such that *M* = *N_b_* × *N* using 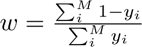

Because of imbalance in the binding labels, we also tried focal loss:

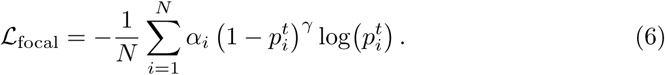

Here 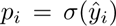 and 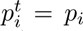 if *y_i_* = 1, else 1 − *p_i_*. The adaptive class weight was calculated *α* = *N ^−^/*(*N* ^+^ + *N ^−^*), applied as *α_i_* = *α* for positive/binding vertices and 1 − *α* for negative/non-binding vertices. Parameter *γ* ≥ 0 is the focusing parameter (typically *γ* = 2). The results from this loss was not significantly different from the adaptive weighted bce loss, so reported models are trained with adaptive-weighted bce.

#### 4.5.2 Partner-specific

The partner-specific model draws on the partner-agnostic case as shown in Figure 1 (c). Here similar to the partner-agnostic model, we have a geometric transformer blocks and a decoding MLP block. The difference lies in the input, output and geometric transformer block. Here, the model takes a pair of coarse protein surface graphs, one belonging to the target protein and another to a potential binding partner. Within the geometric transformer block, we update states of each node on both graphs by first computing individual states, i.e., *s*_0_ = concate(*q*_0_, ***p***_0_ ) and *s*_1_ = concate(*q*_1_, ***p***_1_ ) on each protein (target = 0, binder = 1) using the same self-attention mechanism discussed in section 4.5.1. Since both protein go through the same geometric block, they share the same weights. Then we compute cross-state states *s*_01_ and *s*_10_ by considering a cross-attention block. More specifically, we generate

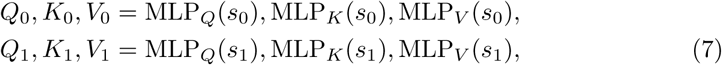

where we have three MLPs for query, key and values and encode *s*_0_ and *s*_1_. To get *q*_01_ and *q*_10_, we compute cross attention scores as

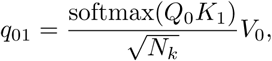

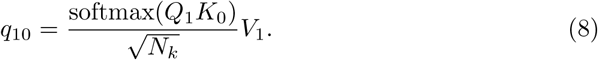

Here, we have kept the dimension of keys equal to *N_k_* for both in the pair. For each protein in the pair, we apply a gated fusion mechanism to update the scalar component of there states *s*_0_ and *s*_1_. For example, for protein 0, we get *q^′^*

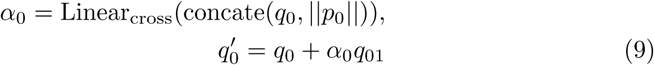

which will then updates states of each nodes in each protein to 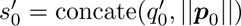 and similarly 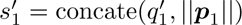.

To train the model, we ask three questions: is node *v_i_* on protein target A a binding site? Similarly is node *u_j_*on protein binder W a binding site? Then are these two vertices make a partner? We perform triplet complementarity loss in that the goal is to place true partner embeddings *antipodal* in the latent space (sim 1), while keeping non-partner pairs far and well-separated from 1. We are essentially flipping the Siamese objective — instead of learning similarity, we learn complementarity, where the “correct” match lives at cosine similarity *→* -1 (antiparallel embeddings).

Taking the embedded per-vertex states 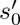 and 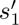 from protein surface A/0 and protein surface W/1, we normalize them to

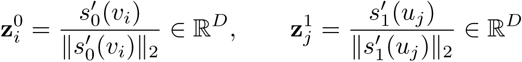

Because they are L2-normalized, the cosine similarity reduces to a dot product:

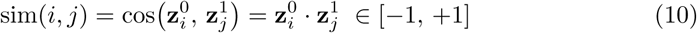

Now, let *P* = *{*(*i, j*)*}* be the set of ground-truth partner pairs, i.e. vertex *v_i_* on A/0 and vertex *u_j_* on W/1 are interface partners. For each positive pair (*i, j*) , we mine *K* hard negatives: the *K* non-partner vertices on W/1 with the most negative cosine similarity to anchor 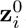, i.e. the non-partners the model currently mistakes as partners.

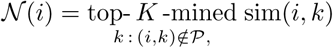

The per-triplet loss then becomes:

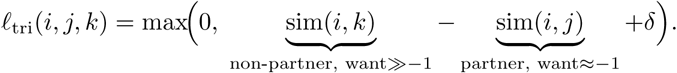

Since cosine similarity spans [-1,+1] with a realistic working gap of ~ 1.0 between a well-trained true partner (sim 1) and a random non-partner (sim ≈ 0), setting *δ* ∈ [0.3, 0.5] demands the model achieve 30-50% of that realistic separation before the loss switches off, strict enough to enforce meaningful antipodal structure, but not so strict that the loss never saturates and destabilizes training. Averaging over all triplets, we get the complementarity/inversion loss:

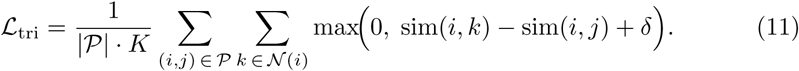

Since we also want for each vertex to independently predict whether it belongs to the binding interface or not, like in the previous section 4.5.1, we calculate 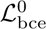 and 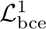, which results into the classification loss of:

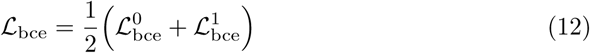

The total training loss is then a regularized sum of these classification losses and complementarity loss in Eq. 11:

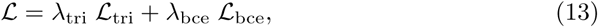

where *λ*_tri_*, λ*_bce_ *>* 0 are regularization parameters, balancing the two objectives. During inference, we get probabilities of binding sites on both the target protein and the partner and from computing Eq.10 we can infer complementarity for all pairs that are *≈ −*1.

### 4.6 Training & evaluation

After performing an exhaustive hyper-parameter search, we use four geometric transformer blocks, and two layers in the decoder MLP. All the MLPs within the geometric transformer blocks have two fully connected layers and two dropout layers with dropout rates of 0.1 for each. We tested neighborhood radius cutoffs of 9Å, 12Å, 20Å and 30Å and found substantial improvements in prediction metrics from 9Å to 20Å but not much from 20Å to 30Å. This is likely because an all node-to-node attention captures some global interaction signatures and refinement from local edge features can be done up to a few amino-acid distance within the neighbor. So, all the results presented below are from models with 20Å radius cutoff for neighborhood attention updating the vector property ***p***. The learning rate is set to 10*^−^*^5^ and Adam Optimizer as implemented in Pytorch Kingma and Ba (2014) was used for training. Trainings were conducted with multiple activation functions such as the Rectified Linear Unit (ReLU), Sigmoid Linear Unit (SiLU), Exponential Linear <u>Un</u>it (ELU) and Gaussian Error Linear Unit (GELU). 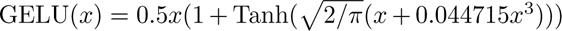 gave the most stable predictions on the test sets, albeit at the cost of slowing the training a bit. The model is trained on batch size of 4 for 1000-1500 epochs or monitored until the validation error did not improve for 20-50 steps set as a patience threshold. Note that this validation set was chosen random’y at 10% of the train sets in Fig. 6.

The hyperparameters used for partner-agnostic model in section 4.5.1 was also used for training the partner-specific model with the addition of cross-attention block that has two heads. Similarly, the MLPs in the cross-attention block have two fully connected layers with intercalated dropout layers of rate 0.1. The model is trained with labels specific to the interface between partners. For inference on two proteins, we can calculate the cosine-similarity matrix and determine binding vertices that have values close to -1 in this matrix to likely be partners. The regularization terms in Eq. 13 are tested in ranges *λ*_tri_ = [0.5, 1] and *λ*_bce_ = [0.5, 1]. The *K* hard negatives were tested at values of 5-10 and *δ* at values of 0.3, 0.4, and 0.5. Model in this text has *K* = 5 and *δ* = 0.4.

The metrics for evaluation of model performance remained the usual metrics used for binary classification such as accuracy (ACC), F1 (the harmonic mean of precision and recall), area under the curve of precision-recall (PR AUC) and the distribution of area under the receiver operating characteristics curve (ROC AUC). F1, PR AUC and ROC AUC are threshold-agnostic metrics as they do not assume a threshold for converting the probabilities to binary labels of 0 and 1. So, most of the results in the paper are reported in these metrics unless they are specified in the text.

## Accessibility

Models will be released on Hugging Face upon publication and will also be available on our SurfGraphPro Github repository with examples for how to run the model on test proteins.

## Software and Data

Training data is curated using PDB IDs of complexes from Dockground database Collins et al. (2022). For benchmarking purposes, the held-out test set include all PDB IDs curated by Gainza et al. (2020) which includes proteins from non-redundant list in PRISM list Baspinar et al. (2014), the ZDock benchmark Vreven et al. (2015), PDBBind Liu et al. (2015) and SabDab Dunbar et al. (2014). Protonation of PDB structure is performed using the Reduce Word et al. (1999) software installed via conda-forge and surface mesh is generated via MSMS Imperiali et al. (1993). Surface mesh cleaning is done via Pymesh Zhou (2018). The 3D surface visualizations are done in PyMOL Schrödinger, LLC (2026).

## Acknowledgments

This work was supported by the Laboratory Directed Research and Development program of Los Alamos National Laboratory under grant numbers 20250639DI, 20250638DI, 20250637DI and 20240734DI. This research used resources provided by the Los Alamos National Laboratory Institutional Computing Program, which is supported by the U.S. Department of Energy National Nuclear Security Administration under Contract No. 89233218CNA000001. A US patent application related to the work is under process. This work is approved for public release by LANL under LA-UR-26-27459.

## Supporting Information

**Supporting Information** The supporting data presents extended analysis of the models and hyperparameter optimization tests.

### 4.7 Core Terminology and Representations

Before describing our approach in detail, we clarify the key geometric representations used throughout this work and their relationships as also illustrated in Fig. 6 (b) and Fig. 1:

#### Surface Mesh

A triangulated representation of a protein’s solvent-excluded surface (SES) generated using MSMS Imperiali et al. (1993). The mesh consists of *vertices* (3D points on the surface) connected by edges forming triangular faces. Each vertex corresponds to a point on the molecular surface at approximately 1 Å resolution.

#### Vertex vs. Node

We distinguish between *mesh vertices* (the fine-grained points in the original triangulated surface) and *graph nodes*, which are the coarse-graphed representations used in our neural network. The raw surface mesh for a typical protein contains thousands of vertices, making direct learning computationally prohibitive.

#### Residue-Centered Patches

To enable efficient learning, we downsample the surface mesh by grouping vertices into *patches* centered on surface-exposed amino acid residues. Each patch comprises all mesh vertices that belong to the solvent-exposed surface of a particular residue. The geometric center of these vertices defines a graph node in our coarse surface representation.

#### Coarse Surface Graph

The final input to the geometric transformer is a graph = ( , ) where nodes *v* represent residue-centered patches and edges *e* connect spatially proximal nodes (within radius *R*). Each node is associated with: (i) ESM-2 embeddings of the corresponding residue, (ii) the patch surface area, and (iii) 3D coordinates of the patch center. This coarse-graphing reduces the problem size from thousands of vertices to hundreds of nodes while preserving essential geometric and chemical information. Figure 1(a) illustrates this multi-scale representation: from atomic structure to fine-grained surface mesh to coarse residue-based graph.

